# Turning Ecology Against Pesticide Resistance: Exploiting Competition in Pest Populations Through Pesticide Use

**DOI:** 10.1101/2023.06.16.545263

**Authors:** Gilberto Muniz-Jr, Maurício Almeida-Gomes, Aliny P. F. Pires, Rafael Dettogni Guariento, Fábio de Oliveira Roque

**Affiliations:** Laboratório de Ecologia, Universidade Federal de Mato Grosso do Sul (UFMS), Brasil; Laboratório de Ecologia e Conservação de Ecossistemas, Universidade do Estado do Rio de Janeiro (UERJ), Brasil

**Keywords:** integrated pest management, mathematical modeling, competition, pest containment, adaptive therapy

## Abstract

Modern agriculture relies on effective pesticides for pest control, but this efficacy is threatened by the evolution of resistance. Although pesticides are novel compounds, pests can quickly develop resistance. Inspired by medical research, we propose an ecologically inspired pest management paradigm that uses pesticides to leverage competitive interactions between pesticide-sensitive and resistant pests. This approach involves reactive pesticide use, monitoring pest responses, and maintaining pest populations below economic injury levels while limiting resistant individuals’ proliferation. A mathematical model shows that managing pests’ abundance at critical levels, rather than aiming for eradication, delays resistant-pest dominance. By reducing resistant individuals’ growth rates and maximizing pesticide-sensitive populations, resistance spread can be hampered. Our findings suggest proactive, ecological, and evolutionary-informed pest management to address threats to food production. This study also highlights opportunities for further research into the ecological aspects of pesticide resistance.

## 1. Introduction

In medical research, understanding how resistance to antibiotics or chemotherapeutics evolves and spread is key to improving treatment strategies, avoiding multi-drug resistance, and minimizing threats to public health (MacLean & San Millan, 2019; Michor *et al*., 2006). Similarly, the food production system, and food security at a global scale, may share similar challenges. Pesticide use to protect crops dates back 4000 years but reports of resistance to pesticides appeared in the literature only at the end of the nineteenth century (Pretty & Bharucha, 2015). However, resistance to pesticides likely arose shortly after the first pesticides were employed. Notoriously, the resistance to pesticides emerges as an evolutionary process, where traits that confer the ability to avoid or tolerate pesticides are selected. Currently, several causes of resistance are known for multiple types of agricultural pests, such as *de novo* mutations, standing variation, or pleiotropic co-option (Hawkins *et al*., 2019).

The growing recognition of widespread resistance to conventional (Hicks *et al*., 2018) and modern (Perez-Jones *et al*., 2007) chemical compounds built for pest control, has led to the notion that conventional management practices must change (Gould *et al*., 2018). However, the recommendation of high doses of pesticide levels to protect crops still prevails (Colin *et al*., 2020), and is based on the assurance of optimized food production through the maximum probability of pest eradication. The use of elevated levels of pesticides can rapidly induce target site selection in favor of resistance genes, producing resistant populations in a few generations (Gould *et al*., 2018), and potentially leading to large economic damages (Hicks *et al*., 2018). Furthermore, through high pesticide levels, current agricultural practices are not only favoring resistance proliferation, but the extinction of many non-target species (Colin *et al*., 2020), contributing to the ongoing, and catastrophic, biodiversity crisis (Díaz *et al*., 2020).

To diminish the development of resistance due to pesticide use, many approaches have been suggested. For example, these include the establishment of spatial pesticide refuges (Takahashi *et al*., 2017), rotation and mixture of pesticides (Cloyd, 2010), optimization of concentration levels based on pest physiological responses (Colin *et al*., 2020), or integrated pest management (IPM), that promotes combined, often non-chemical, solutions to pest control (Peterson *et al*., 2018). Many of these proposals have been implemented at large scales, but recent empirical data has shown that such strategies are not preventing the evolution of resistance from emerging (Hicks *et al*., 2018). In addition, even the straightforward recommendation for a reduction in dose rates can produce an opposite outcome by hastening resistance through the gradual gathering of mutations (Muniz-Junior *et al*., 2023). Therefore, considering the widespread resistance to pesticides, the effectiveness of current pest management practices seems transient, and a better strategy would be to use lower pesticide levels that control pest populations, but also prevent the prevalence of resistant pests, building agricultural systems that meet both production and environmental goals.

Inspired by recent advances in medical research, where curative methods seek to primarily control the spread of resistant populations (Hansen & Read, 2020a), and the discovery of new drugs for therapeutic use is heavily constrained (Pui & Evans, 2013), we aimed to explore how a reactive use of pesticides can delay the ecological dominance of resistant pests and also constrain pest abundance (i.e. pest containment) at levels in commitment with economic, ecological and evolutionary aspects. Using a classical mathematical model of species competition, we explore the dynamics of pesticide-resistant and sensitive populations under different scenarios of pesticide use, resistance prevalence, and tolerable pests’ population sizes. Our simulations and theoretical predictions show that a reactive use of pesticides supports the co-existence of pesticide-sensitive and resistant individuals in the pest population, diminishing the growth rate of resistant populations, and ultimately delaying the time of management failure due to resistance.

## 2. Materials and Methods

### 2.1. From medical therapy to crop management: the concept of pest containment

Our proposal of using pesticides to promote pest containment and delay resistance dominance is inspired by recent empirical and theoretical advances in cancer (Gatenby, 2009), and infectious diseases (Hansen *et al*., 2020; Letten *et al*., 2021) research, where the therapeutic use of chemotherapeutics or antibiotics is compromised by the proliferation of resistant cells. This new paradigm in treatment protocols, known as adaptive therapy, emerged to focus on limiting the proliferation of resistant individual cells, instead of immediate cure (Hansen & Read, 2020a). We present a similar theoretical foundation, but with assumptions closer to the environmental and agricultural contexts. Instead of using a fixed schedule, pesticide doses and usage are based on how pests respond to pesticides, it is, therefore, reactive. This contrasts with conventional pesticide use, which uses predetermined doses based on pest mortality curves (Krieger, 2010). Second, it is designed to leverage competition between sensitive and resistant individuals to improve resistant population control. We assume spatial homogeneity and that resistance is widespread, already present in individuals of a potential infestation, or very likely to emerge (Bras *et al*., 2022). Rather than attempting to eradicate pests, the goal is to keep pest population size at certain levels, maintaining sensitive individuals in order to competitively suppress resistant ones. Such a goal is defined here as pest containment, a term that was borrowed from cancer research, where tumor containment may be desired when a cure is rarely achievable (Hansen *et al*., 2017).

### 2.2. Model formulation and simulation

We built a model with two types of competing populations, sensitive and fully resistant to pesticides, with population sizes *S(t)* and *R(t)*, respectively. The initial values for both populations are *S*0 and *R*0, and the total pest population size is denoted by *N(t) = S(t) + R(t)*, leading to *N* 0 = *S*0 + *R*0. Our model formulation does incorporate mutation processes and complex breeding/inheritance systems of distinct pests (Helps *et al*., 2017). However, based on our assumptions, and the essential features to contrast the effects of pesticide dosing on the dynamics of resistant and sensitive populations, our result directly sheds light on how competitive exclusion will be affected by pesticide dosing schemes. Our goal is to contrast the effects of pesticide dosing on the dynamics of resistant and sensitive populations, highlighting on how pesticide dosing schemes affect the rate at which resistant individuals outcompete sensitive individuals. The population size N0 is also the target size of the pest population where it must be contained. We define it as the critical pest population size (CPPS). In other words, a pest population size when the pesticide intervention should start. Ideally, up to the CPPS pest abundance should not cause significant crop yield losses, or should produce tolerable losses compared to the management costs. Pest dynamics are described by a modified Lotka-Volterra competition model, and the equations describing the population dynamics are given by:

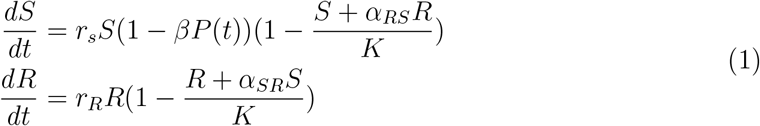

Where 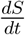 and 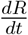 denote derivatives, *r*_*S*_ and *r*_*R*_ are *per capita* growth rates; the quantity *P* (*t*) is the pesticide dose at time t, and *β* represents a constant that translates dosage into a reduction in pest population growth rates. *K* is the pest population carrying capacity (shared by both sensitive and resistant populations), *α*_*ij*_ is the competition coefficient that determines the effect of population *i* on *j*.

The model was parameterized with values for pest growth rates within the range observed for weeds and fungi (Meyer *et al*., 2021; Oreja *et al*., 2021; Suberkropp *et al*., 2020). However, we also provide analytical solutions to our results, diminishing the reliance of our conclusions, at least qualitatively, on specific parameter values used for the simulations, and therefore, on any specific pest model system. The parameter values used for the simulations are provided in Table 1.

**Table 1.** Parameters description and values used in the numerical simulations of the model.

| Parameter | Description | Value | Units |
| --- | --- | --- | --- |
| $r_S$ | Sensitive population proliferation rate | 0.015 | $\frac{1}{d}$ |
| $r_R$ | Resistant population proliferation rate | 0.015* | $\frac{1}{d}$ |
| $P_{max}$ | Maximum pesticide rate | 1.5 | $\frac{ppm}{d}$ |
| $\beta$ | Dose-response pesticide fitness reduction on pest population | 1 | $\frac{1}{ppm}$ |
| $\alpha$ | Competition coefficients among pest populations | 1** | - |
| K | Populations carrying capacity | 1 | $\frac{mg}{cm^2}$ |
| $R_{frac}$ | Initial resistant population % fraction | 0.1 - 1 % | - |

\* Distinct intrinsic growth rates for the resistant population are evaluated in the Appendix S1
\*\* For the results presented in figure 3, $\alpha_{RS}$ was kept constant and $\alpha_{SR}$ varied accordingly.

#### Our analysis is built based on the following assumptions

(i) The growth rate of sensitive individuals is positive in the absence of pesticide use, and decreases as pesticide doses are increased, becoming negative at *P* = *P*_*Max*_; (ii) Competition coefficients can be ignored if pest composition is restricted to a single species because the effect of S on R and vice-versa would be equivalent. However, we also explore a scenario where sensitive and resistant individuals present asymmetrical competition effects; (iii) Mutations that differentiate sensitive individuals into resistant ones are neglected. Therefore, resistance proliferation is primarily driven by preexisting resistant individuals. As the main outcome of our simulations (i.e., our output criteria) we will use the time to management failure (TMF). We define the TMF as the time until the pest population exceeds the CPPS level. We also defined a criterion that management failure occurs when the pest population size is 20% larger than CPPS. This 20% buffer would make sense in practical terms, due to imperfect pest abundance estimations. However, smaller buffer values, or even the lack of this buffer, does not qualitatively affect our conclusions or interpretations.

We also evaluated a scenario where resistance to pesticides implicates fitness costs to the resistant populations through a reduction in growth rates (Kliot & Ghanim, 2012). This would make resistant and sensitive populations diverge about their ecological equivalence, beyond a pesticide-induced reduction in growth rates. These results are presented in the Appendix S1 and no qualitative differences were observed with such a feature added to our model. In fact, a fitness cost on resistant populations only intensified the desired effect of reactive pesticide use.

We considered as pesticide use protocols the intensive pesticide use, **IPU**: *P* (*t*) is fixed at *P*_*Max*_ for every *t*; and the **reactive pesticide use**: withdraws pesticide administration, *P* (*t*) is set to zero, once a reduction threshold (i.e., a certain proportion decrease from the limiting CPPS *N* 0) is achieved and reinstates it to *P*_*Max*_ once the pest grows to the CPPS. As reduction threshold levels for our simulations, we used 25% and 50% of the CPPS, but an analytical generalization of the effect of the magnitude of these reduction levels is provided in the Results section.

### 2.3. Numerical solutions

Equation solutions were obtained using the class scipy.integrate.ode for the Python programming language. The reactive protocol was implemented as a sequence of intervals of length Δ*t*, with *P* = *P*_*Max*_, where *P* is controlled between intervals according to the proposed protocol of pesticide use. All simulations were carried out in Python 3.9, using the modules Scipy = 1.4 and Numpy = 1.16. Codes for reproducing the simulations are available at: https://github.com/rafaelguariento/reactive_pesticides_use

## 3. Results

### 3.1 The role of pest population size and reduction thresholds on pest containment

Throughout the results, we compare the model-predicted time to management failure (TMF) for both intensive (*TMF*_*IPU*_ ) and reactive (*TMF*_*R*_) pesticide use protocols. We observed that *TMF*_*R*_ is progressively greater than *TMF*_*IPU*_ as pests are contained at higher levels of CPPS and closer to such density (i.e., with smaller reduction thresholds) (Fig. 1). This result reflects the nature of the reactive design approach, which is to leverage the competition between sensitive (*S*) and resistant (*R*) populations to reduce the growth rate of resistant individuals. As a way to demonstrate the role of competition, the variation of *R* (i.e., *R*(*t* + *dt*)) for a small arbitrary time interval, *dt*_*R*_, can be approximately related to the instantaneous variation in *R* (i.e., *dR/dt*). Therefore, based on Eq. 1 *dR/dt*_*R*_ would be equal to: 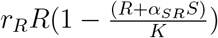. Rearranging the former equation to isolate *dt*_*R*_, we have:

**Fig. 1.**
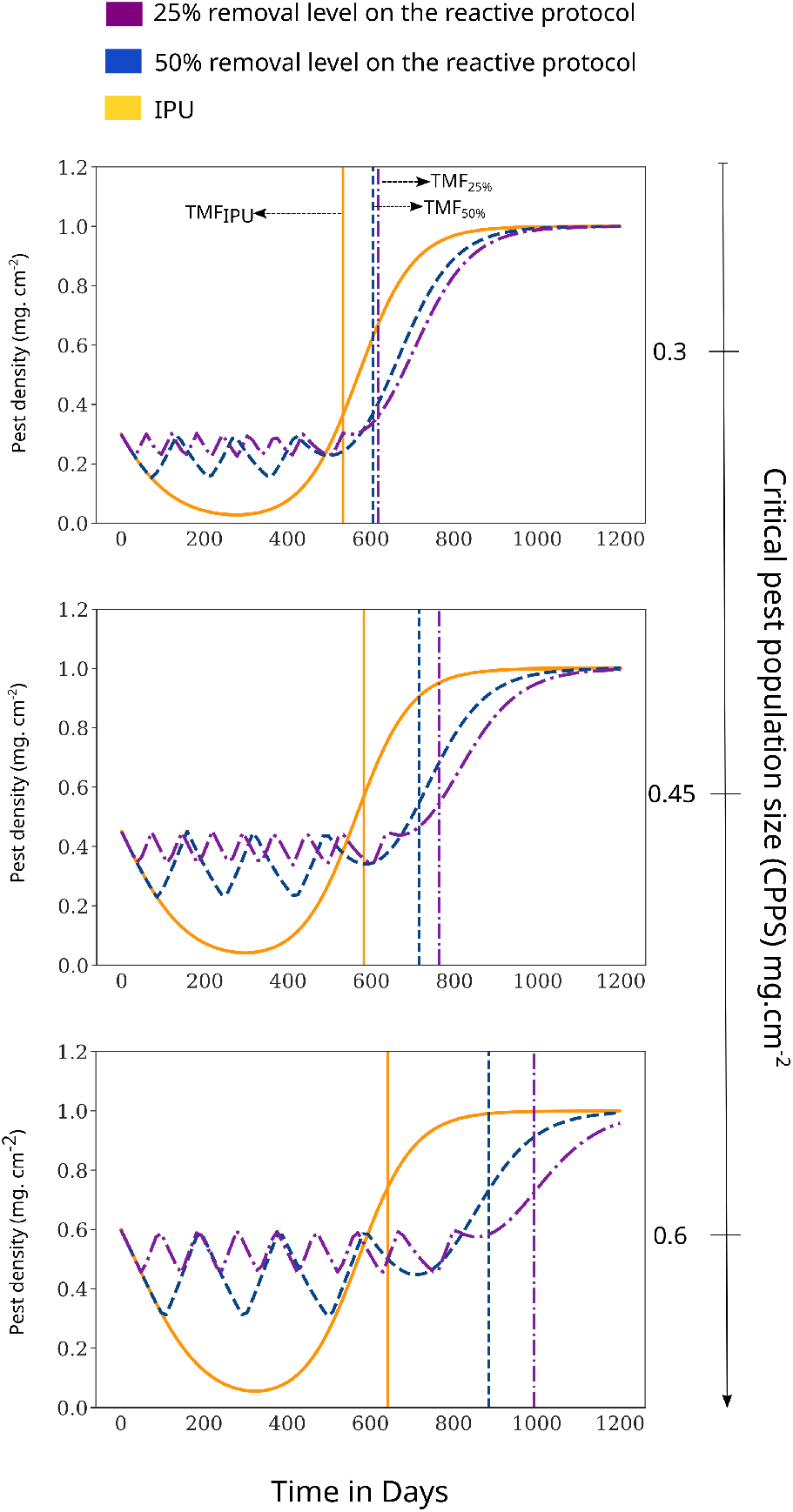
Dynamics of total pest abundance (*N* (*t*)) for simulations under both IPU and the reactive pesticide protocols for pest populations at different target densities (i.e., Critical pest population size - CPPS). Vertical lines indicate the TMF for the IPU (yellow) and reactive protocols with 50% (blue) and 25% (purple) removal reduction thresholds, respectively

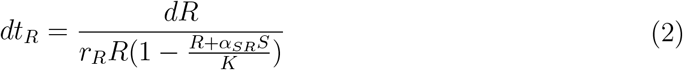

From Eq. 2, greater (*S*) implies a smaller denominator and hence higher *dt*_*R*_. We can assume that the density of S will be kept greater under the reactive protocol, compared to the intensive pesticide use, because it allows sensitive individuals to grow due to pesticide withdrawal. Thus, based on Eq. 2, and the preceding assumption, *dt*_*R*_ |_*Reactive*_ will be greater than *dt*_*R*_ |_*IPU*_ . In other words, the time it takes for the resistant population to grow to an arbitrary size is maximized by the reactive protocol compared to the intensive use, which translates into reduced pesticide-resistant population growth rates.

The performance of the reactive approach, compared to the IPU, increases with the CPPS as sensitive individuals can be kept at higher densities for longer (Fig. 2). This implies that a better reactive design strategy would have to promote a minimal reduction level and maintain the pest at the limit of the CPPS as closely as possible (i.e., optimal containment), in order to maximize competition. Considering such an assertion, to promote a minimal reduction and maintain pests at the CPPS limit, it would require *P* doses to keep *dN/dt* = 0. The rate *dN/dt* = *dS/dt* + *dR/dt*, and solving the system of equations for *P* as in Eq. 1 when *dN/dt* = 0, we get:

**Fig. 2.**
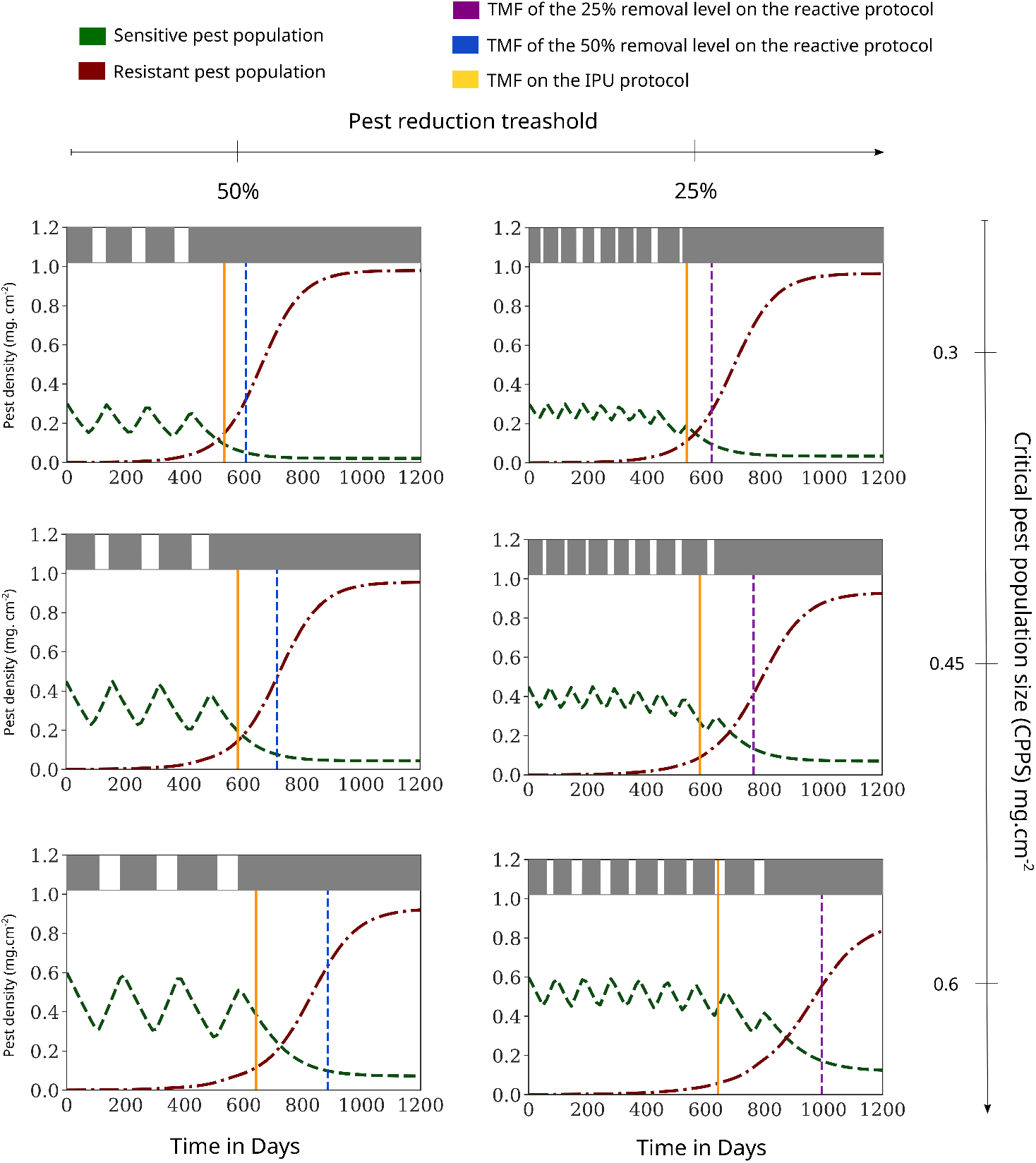
Dynamics of individual populations of sensitive (green) and resistant (red) pests, for simulations under the reactive pesticide protocols with 50% (left) and 25% (right) removal reductions thresholds, for pest populations at different target densities (i.e., Critical pest population size - CPPS). Vertical dashed lines indicate the TMF of IPU (yellow) and reactive protocols with 50% (blue) and 25% (purple) removal reduction thresholds, respectively. The top bar indicates when pesticides are being applied (gray), or withdrawn (white), according to the respective pesticide use protocol

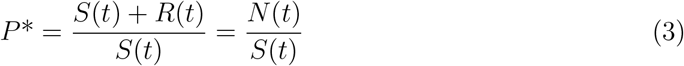

Any value of *P > P* * will lead to *dN/dt <* 0. The quantity *P* * will always be lower than *P*_*Max*_ because, by definition, *P*_*Max*_ leads to negative values of *dN/dt*. Thus, the reactive protocol would also be characterized as less intensive because it would require lower P doses than the standard protocol to manage pests. However, based on Eq. 3, this property wears out as *N* (*t*) increases, which translates into higher *P* * rates as the greater the CPPS, or as closer, the containment is to the CPPS. This last feature can be visualized in Fig. 2 as the periods when pesticide administration is withdrawn become shorter for the 25% removal threshold, compared to the 50% threshold.

### 3.2. Pre-existing resistance levels and competition strength effects on containment

To better visualize the performance of the reactive protocol compared to the IPU, we calculated the Log Response Ratio (LRR) of the values of *TMF*_*R*_ and *TMF*_*IPU*_ across different values of the initial prevalence of resistant individuals in the pest population and the competition effect that sensitive individuals would exert on resistant types (Fig. 3). We observed that the greater the prevalence of resistant individuals, the lower the benefit of the reactive protocol. However, the performance of the reactive protocol increases as the competition effect of sensitive individuals on resistant types increases. As shown in Eq. 2, the larger the competition coefficient *α*_*SR*_ and the abundance of sensitive individuals, the larger the negative component in the denominator, contributing to the increase in *dt*_*R*_. Analogously, an increase in the prevalence of resistant individuals would reduce the contribution of *S*, and its negative effect on the growth rate of the resistant population.

**Figure 3.**
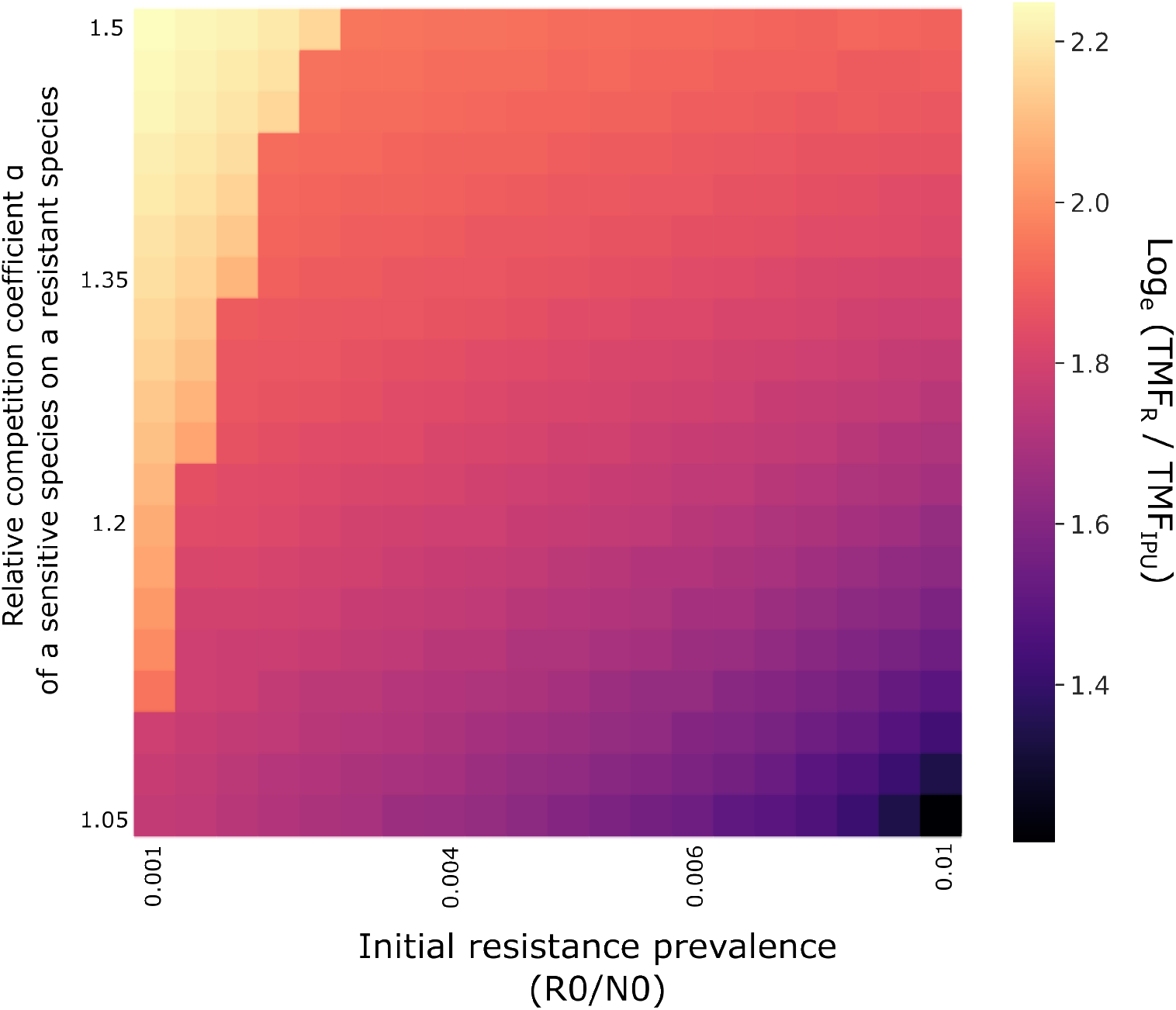
Log response ratios (LRR) of the performance of the reactive protocol, in terms of the time to management failure (TMF), compared to the intensive pesticide use (IPU). The LRR is computed for distinct relative competition effects of the sensitive population on the resistant population (*α*_*SR*_) and values of initial resistant prevalence in the pest population. We fixed the values of *α*_*RS*_ and varied the values of *α*_*SR*_ to achieve a variation in the relative competition coefficients among populations. For the simulation of the reactive protocol, we used a removal level = 50%

## 4. Discussion

Under a scenario where resistance to pesticides is already present in crops, we showed that the time until management failure, due to the dominance of pesticide-resistant pests, can be prolonged when pesticide use exploits competitive interactions among individuals, compared to the standard intensive use. Ultimately, this result diminishes the reliance of the success of pest management in the discovery of novel pesticides, and builds a framework for an ecologically-informed pesticide use protocol. We showed that the performance of the pesticide reactive use depends on the initial fraction of resistant individuals and the ability to hold the pest proliferation at a relatively large population size, ideally close to the limit of economic injury levels. In the following, we discuss how these results relate to the underlying ecological mechanisms and the opportunities for a more efficient and sustainable approach to pesticide use.

### 4.1. The Ecology of Resistance Proliferation and Suppression

Throughout the years, agriculture faced many ecological and evolutionary challenges, that promoted a plethora of strategies to minimize resistance evolution, such as high-dose-refuge strategies, culture rotation and administration of multiple control methods (REX consortium, 2010). However, management practices should also be able to effectively control resistant populations once resistance has been established in the population. We can explain the benefit of a reactive use of pesticides, compared to standard intensive use, in minimizing resistant population growth asking when it would be beneficial to maintain sensitive populations to suppress resistant individuals, and when it might be better to maximize the tolerable pest density. Both propositions would preclude resistant individuals from competitive release. As long pest population growth is governed by the growth of the resistant population, the more a pesticide dosing protocol can reduce resistant population growth rate, but maintaining overall pest population size below desired levels, the longer the containment of population pest size and management success. From our theoretical analysis, irrespective of the intrinsic growth rates of sensitive and resistant populations, the larger the sensitive population size, the lower the resistant population growth rate. Comparable results were obtained in a study of antibiotic resistance in bacteria (Hansen *et al*., 2017), a result that is driven essentially by the competition among the interacting sensitive and resistant populations. If we compare the sensitive population size under IPU and the reactive approach, the population size of sensitive individuals will be always greater under the reactive approach because it provides an opportunity for the sensitive population to recover. Therefore, the reactive approach will better exploit the competition among sensitive and resistant populations in suppressing resistance expansion. This implies that, under our assumptions, the reactive use of pesticides will always be superior to the intensive use in delaying management failure, ascertaining that such an approach can be successful under a broad array of conditions. As highlighted in studies with cancerous cells, even when different growth models are used, such a conclusion would remain (Viossat & Noble, 2021). We also observed that the benefit of reactive pesticide use is stronger as the CPPS increases. This could be explained simply because sensitive populations can be kept for longer, indicating that the most effective management would maintain the pest population size as large as possible, maximizing competition. Consequently, a crucial step in dealing with resistance that is already present in the pest population is the recognition of a critical pest population size (i.e., CPPS). While interventions at threshold densities have been proposed under an integrated pest management (IPM) approach (Green *et al*., 2020), they are not always applicable. In fact, in many cases, thresholds have not been established for many pests (Barzman *et al*., 2015). On the other hand, IPM emerged in the field of insect pest control, where the use of intervention thresholds was successful (Peterson *et al*., 2018). Such threshold densities are viewed as practical rules, used to determine when to take management action. They become predictions of when a pest population is going to reach densities that will cause yield losses comparable to the insect management costs, known as the economic injury levels (EIL). Therefore, it is pivotal that pest management should be designed to keep the populations below EIL, to avoid severe short-term economic yield losses. However, as stressed in this study, it is also important to take into consideration possibilities for the effective use of pesticides along with the establishment of threshold densities, particularly when considering pesticide resistance evolution (Green *et al*., 2020). Although there have been efforts to define economic thresholds for weeds (Keller *et al*., 2014), for example, obtaining such estimates is challenging because they usually appear as a community of multiple species and have a long-term impact through a persistent seed bank (Sattin *et al*., 1992). Therefore, it may become difficult to establish scientifically sound pest economic injury levels for all major pests in all major crop varieties and cultivation environments. As our study suggests, whether it is best to approach pests with intensive pesticide use or to adopt a containment strategy will depend on the feasibility of threshold-based decisions.

### 4.2. Taking widespread resistance into consideration

Our proposed strategy works as an alternative management choice when resistance is unavoidable. Although this condition may seem uncommon, evolutionary and ecological mechanisms show that this might be common. The low diversity of intensive monocultures of crop species and genotypes are common features of modern agriculture (Martin *et al*., 2019), and such a feature exerts strong directional selection on pests to overwhelm control measures. In addition, it was recently highlighted that the evolutionary history of a species might affect the likelihood of developing resistance, making some species more predisposed to evolve resistance than others (Walsh *et al*., 2022). In this context, phytophagous arthropods, for example, have been suggested to be preadapted to handle chemical insecticides since they have evolved to tolerate and resist their host plant allelochemicals (Després *et al*., 2007), irrespective of prior exposure to pesticides. Such cross-resistance, as resulting from prior exposure to a different toxin in a species’ evolutionary history, can make resistance more widespread than previously thought (Bras *et al*., 2022). Pests may be pre-adapted to pesticides due to exposure to natural inhibitors such as host defense compounds or pathogen toxins, or as a pleiotropic effect of traits unrelated to chemical resistance, whereas standing variation may accumulate through neutral processes and support pesticide resistance (Hawkins *et al*., 2019). When assessing the risk of a pest evolving to overcome a control measure, it is important to consider various sources of resistance. For example, a requirement for multiple mutations or complex resistance mechanisms that are unlikely to arise *de novo* does not exclude resistance risk from the interspecific transfer (Hedrick, 2013). Therefore, the multiple sources of resistance before any control measure is deployed would increase the risk of control failure for novel compounds. In the long-term, the ability to deal with resistant pests to any know pesticide only using a different scheme of pesticide use would contribute to the resilience of the food production system in the face of technological shortages, and diminish the risk imposed by such latent threats. Our model results also point out that the effectiveness of the reactive strategy should improve as lower is the proportion of the resistant population in the total population. Thus, it is pivotal that management practices should be able to effectively control resistant populations once resistance establishes in the pest population.

### 4.3. Challenges and Opportunities of the Reactive Use of Pesticides in Pest Management Strategies

On the social spectrum, arguing to end-users that total eradication of pests is not needed to assure their economic returns will require a major shift in agriculture (Pilcher & Rajotte, 2008). Applying insecticides at levels needed to reduce the target pests’ fitness to zero has been proposed to help manage insecticide resistance (Gardner *et al*., 1998). However, despite the fact that this practice may reduce the likelihood of the selection of single-site mutations, it may increase the proliferation of resistance due to the accumulation of resistance factors (Muniz-Junior *et al*., 2023). We showed that the reactive use of pesticides can hold the proliferation of resistance using lower dosing of pesticides compared to a standard protocol of IPU. Therefore, fulfilling an important aspect of sustainable pesticide use (Pretty & Bharucha, 2015). We also showed that the larger the threshold for pest removal, the lower the total use of pesticides throughout the management. This is because pesticides are withdrawn for longer periods, so the pest population can recover until the pest reaches CPPS. On the other hand, we observed that the effectiveness of the containment (i.e., the time until the TMF) decreases as large as the threshold pest removal. In other words, an idealized containment (i.e., the best containment possible) would require greater dosing of pesticides than schemes of pesticide use that are turned off for greater periods. This suggests that, as similarly stated for the establishment of critical pest population sizes, understanding the allowed TMF for different crops, would further improve the potentially less intensive use of pesticides under a reactive scheme. For example, different harvesting strategies, such as early harvesting before pest propagation (Ramalho, 1994), have been used to suppress the proliferation of resistance. The ability to manage the harvesting time within a workable timeframe makes the reactive use of pesticides especially interesting and unique when compared with the adaptive approach of chemotherapeutics or antibiotics used in medical applications. As long as crops do not need to be maintained for an indeterminate period, but will be harvested at a specific time, the reactive dosing scheme does not need to hold the pest population for as long as possible, but for the time until harvesting is needed. Therefore, pesticide use schemes that do not lead to the greatest TMF possible, might be appropriate for different types of infestation or different crops, minimizing the use of pesticides and resistance proliferation, but ensuring the crop return. Instead of exploiting the competitive interaction within a single population (i.e., intraspecific competition between resistant and sensitive individuals), the reactive use of pesticides can also benefit from the interspecific competition that may happen under multiple infestations. Crops species interact with multiple organisms throughout their life cycle. As a result, they are commonly attacked by multiple species of pests (Magalhães *et al*., 2018). Our results suggest that the performance of the reactive use of pesticides would increases in multiple infestations if one of the pest species is still susceptible to the pesticide and is also the best competitor. This feature provides an additional technological avenue for pest containment, as multiple infestations can be taken as an opportunity to minimize resistance proliferation if pest species differ in their likelihood to present resistance to pesticides (Zhao *et al*., 2017), and also differ in their competitive abilities. This aspect also makes the reactive use of pesticides unique compared to the analogous approach in medical applications, as the likelihood of infestations by multiple species is particularly relevant only in crop ecosystems. The significance of local interactions in influencing the dynamics and structure of communities and populations has been a prominent focus of ecological research in recent decades (Chase & Leibold, 2009), helping us to predict, for example, pesticide use and contaminant effects on communities and ecosystems (Rohr *et al*., 2006; Zhao *et al*., 2017). How such local interactions might shape the ability of individual variance to thrive, has become a flourishing research avenue in medical research (Martincorena & Campbell, 2015; Martincorena *et al*., 2017).In this context, we demonstrate here that these ecological interactions could offer novel insights into addressing the evolution of pesticide resistance.

As theory progresses, both laboratory experiments and human and animal trials have been conducted for testing the efficacy of antibiotics and chemotherapeutics administration under the same principles that we modeled for pesticide use. The results, so far, confirm that such an approach delays the dominance of resistance (Brady-Nicholls & Enderling, 2022; Hansen & Read, 2020b; Wang *et al*., 2021) and substantiates the predicted mechanisms that lead to the observed patterns (Strobl *et al*., 2021). In food production systems, high-detail fine-tuning pesticide delivery and monitoring for classification and density estimation of pests already exist (Tetila *et al*., 2020; Zhang *et al*., 2021), which would facilitate the establishment of CPPS levels and a reactive pesticide administration in terms of pest response. Fitting model expectations to response data from crops under distinct management protocols would serve to test hypotheses related to the mechanisms that reduce the growth of resistant individuals under a reactive approach. For example, as shown by recent studies in medical research, modulation of the goodness-of-fit between empirical data and model dynamics expectations under specific model formulations (e.g., with fitness costs associated with acquiring resistance or with distinct threshold reductions), would indicate the importance of variables or efficacy of certain protocols in explaining management dynamics and the potential drivers of the reactive approach success (Hansen & Read, 2020b; Strobl *et al*., 2021). Thus, with the increasing availability of response data and dynamics of crops under a reactive pesticide administration, such an approach can be more carefully customized to crop and pest types who stand to benefit from it the most.

## 5. Conclusion

Here, we highlight the opportunity to reconcile ecological and management interests with a novel theoretical approach. The reactive use of pesticides shares many similarities with the medical application of antibiotics and chemotherapeutics (Hansen *et al*., 2020, 2017; Letten *et al*., 2021), but novel opportunities and challenges of the reactive approach in food production systems make such an approach unique for this setting. Such uniqueness arises from the exploitation of competitive interactions within and among multiple species, the establishment of economic injury levels of different pest densities, and trade-offs between the effectiveness and the environmental benefits of intensive vs less intensive (reactive) pesticide use. In addition, we foresee interesting avenues of further theoretical development of the reactive use of pesticides such as the ones observed in medical applications, such as exploring distinct growth models (Viossat & Noble, 2021), continuous development of resistance through mutations (Hansen *et al*., 2017), and the role of multiple pesticides in managing pests and their application through a reactive use (West *et al*., 2019). Thus, we recommend and strongly hope that both academics and decision-makers follow lessons from non-conventional pest management practices, and actively engage in the evaluation of evolutionary and ecological consequences in the development and employment of new methods. With increasing awareness that pesticide-resistant organisms are set within complex species interactions, it would be short-sighted not to take advantage of a wealth of helpful theory and models at the frontier of ecological research.

## 6. Acknowledgments

We thank CAPES, CNPq, and Fundect for continuous support through research grants.

