## Appendix S1 for "Turning Ecology Against Pesticide Resistance: Exploiting Competition in Pest Populations Through Pesticide Use"

### Turning Ecology Against Pesticide Resistance: Exploiting Competition in Pest Populations Through Pesticide Use

Gilberto Muniz-Jr[1,*] , Maurício de Almeida-Gomes[1], Aliny P. F. Pires[2], Rafael Dettogni Guariento[1] and Fábio de Oliveira Roque[1]

1. Laboratório de Ecologia, Universidade Federal de Mato Grosso do Sul (UFMS), Brasil;

2. Laboratório de Ecologia e Conservação de Ecossistemas, Universidade do Estado do Rio de Janeiro (UERJ), Brasil;

**Figure A1** - Dynamics of total pest abundance (N(t)) for simulations under both IPU and the reactive pesticide protocols for pest populations at different target densities (i.e., Critical pest population size - CPPS). Vertical lines indicate the TMF for the IPU (yellow) and reactive protocols with 50% (blue) and 25% (purple) removal reduction thresholds, respectively. On the left, are the results for equal growth rates among pesticide-sensitive and resistant populations. On the right, the results when the resist populations have 25% reduction in growth rates compared to the sensitive population due to the fitness cost of acquiring pesticide resistance.

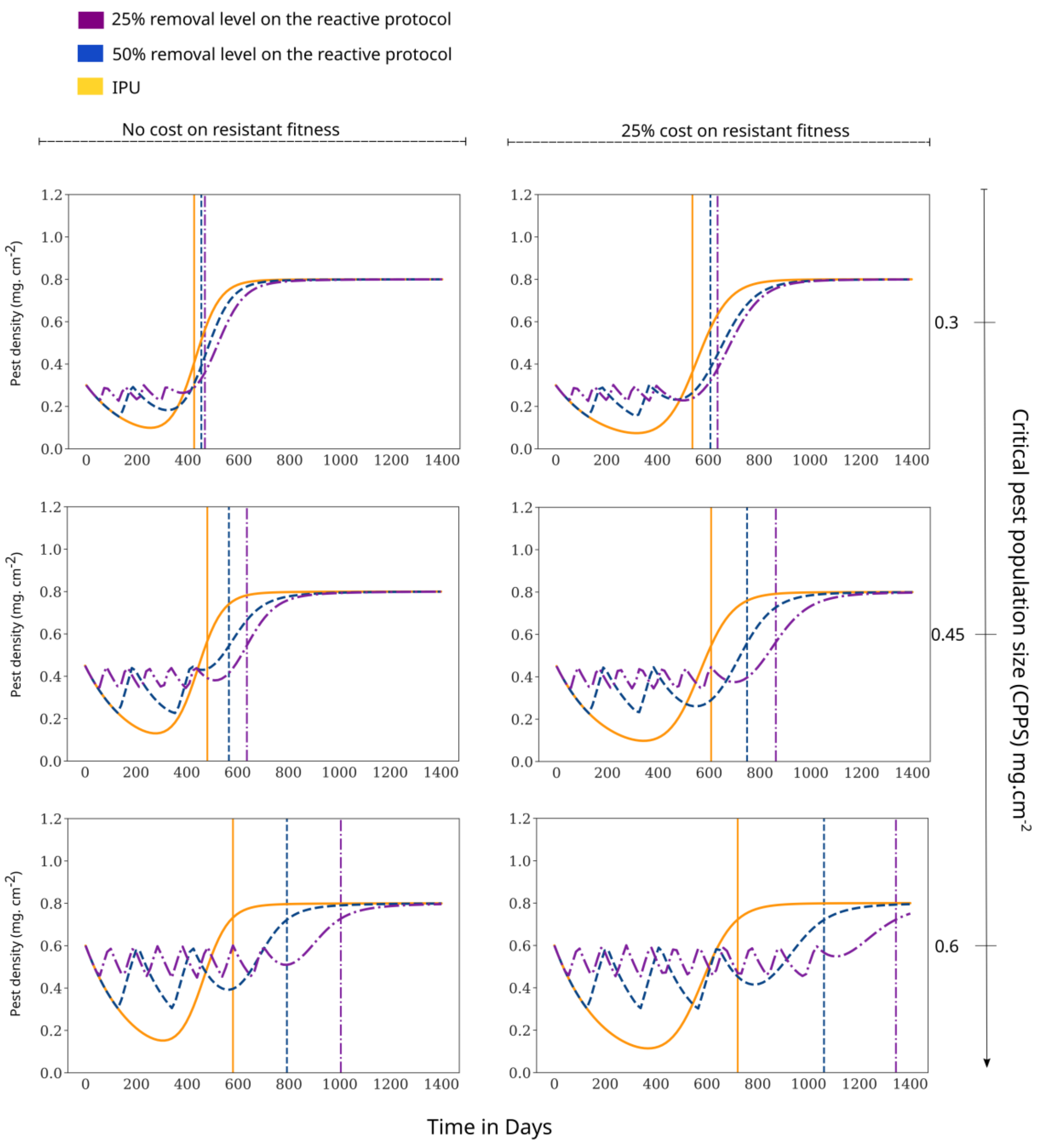
